# Machine learning differentiates between bulk and pseudo-bulk RNA-seq datasets

**DOI:** 10.1101/2025.06.27.661895

**Authors:** Boon How Low, Md Mamunur Rashid, Kumar Selvarajoo

**Author notes:** **Correspondence.**; (Kumar Selvarajoo). Equal contributions (Boon How Low, Md Mamunur Rashid).

## Abstract

Modern synthetic data generators and deconvolution methods rely heavily on single-cell (sc) RNA-seq data. Aggregated scRNA-seq (pseudo-bulk) is commonly assumed to closely match true bulk RNA-seq, making it a dependable benchmark for developing and evaluating new bioinformatics methods. Here, we investigated paired bulk and scRNA-seq datasets using machine learning techniques to assess the fidelity of pseudo-bulk profiles. Our results demonstrate that pseudo-bulks differ substantially from bulk RNA-seq in both analytic metrics and biological processes.

## Main

RNA sequencing (RNA-seq) has revolutionized our ability to characterize gene expression landscapes in health and disease [1, 2]. While bulk RNA-seq provides tissue-level biological insights, single-cell RNA-seq (scRNA-seq) enables the resolution of cellular heterogeneity within complex tissues [3]. Despite the transformative impact of scRNA-seq in advancing our understanding of complex biological systems, its application to individual patients in biological or pathological processes remains time-consuming, costly, and difficult to scale. Moreover, scRNA-seq data are inherently sparse due to technical and biological dropout events, limiting the recovery of low-abundance transcripts and complicating downstream analysis.

The recent developments of synthetic data generators aim to recover missing signals and model disease progression [4]. However, they often rely on pseudo-bulk (PB) profiles, generated by aggregating scRNA-seq data across cell populations, as reference standards for validating synthetic datasets. PB strategies have been widely used to bridge scRNA-seq and bulk transcriptome, serving as a practical proxy for true bulk RNA-seq in various algorithms [5,6], including cell type deconvolution, differential expression analysis, and synthetic data benchmarking. However, the fundamental assumption that PB profiles faithfully recapitulate true bulk RNA-seq signals remains unexplored. This knowledge gap is particularly concerning given the widespread use of PB-based approaches in computational biology [7].

To evaluate the fidelity of PB profiles, we present a systematic comparison using matched PB and bulk RNA-seq datasets sourced from three independent studies: breast cancer with clinical subtypes (GSE176078) [8], high-grade serous ovarian cancer (GSE217517) [9], and cell lines derived from induced pluripotent stem cells (iPSC; GSE226163) [10]. We preprocessed each dataset uniformly using standard data quality control pipelines for downstream analyses (Methods). PB profiles were generated by aggregating scRNA-seq counts per sample. To adjust for sequencing depth/library size differences, raw counts were normalized to counts-per-million (CPM), and log-transformed with a pseudo-count of 1. Genes expressed in < 20% of samples were excluded to ensure statistical robustness. To first assess the global expression patterns of PB and bulk RNA-seq samples, we applied

Sparse Principal Component Analysis (SPCA) on 0-centered expression matrices and compared sample distributions along the first two principal components (PCs). In all three datasets, PC1, accounting for the largest proportion of variance, clearly separated PB and bulk transcriptomes into distinct clusters. This indicates that differences in data modality are the primary drivers of transcriptomic variation (**Figure 1a, Supplementary Figure 1**). In contrast, PC2 captured biologically meaningful heterogeneity, including clinical subtypes in breast cancer, intra-tumor variability in ovarian cancer, and distinct cell line identities in the iPSC dataset (**Supplementary Figure 2a**). While these results suggest that PB profiles retain important biological signals of the bulk (PC2), the overall transcriptomic landscape is largely influenced by inherent differences between bulk and single-cell-derived data, reflecting distinct statistical and biological characteristics of each modality (PC1).

**Figure 1.**
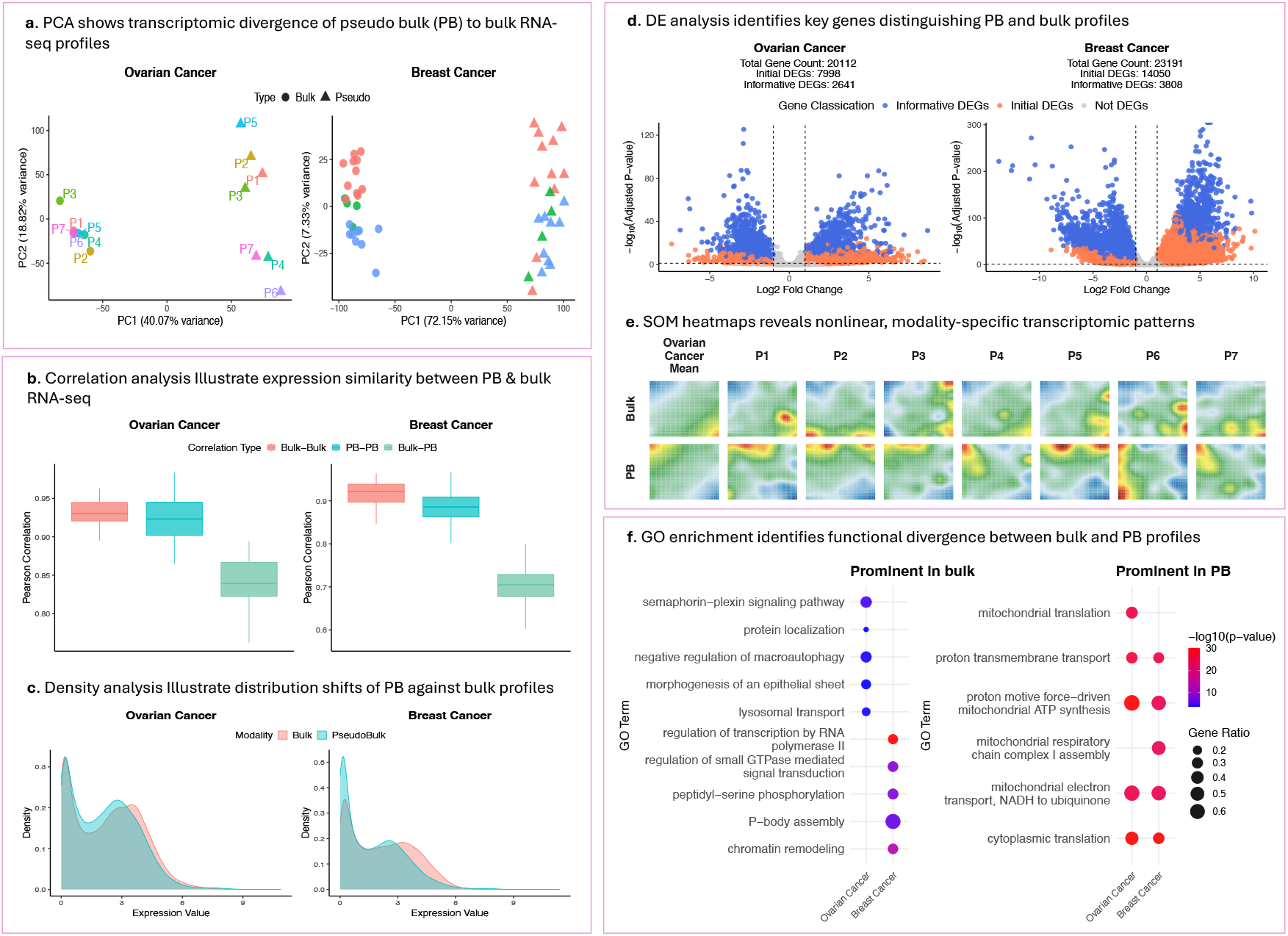
Systematic transcriptomic divergence between pseudo-bulk (PB) and bulk RNA-seq profiles. **a**. SPCA shows that PC1 distinctly separated samples by data modality, with underlying sample heterogeneity captured along PC2. **b**. Pearson correlation analysis shows consistently lower similarity between PB and bulk profiles compared to within-modality (bulk-bulk and PB-PB) comparisons. **c**. Density analysis highlights a right-shifted expression distribution in bulk samples, indicating a greater proportion of highly expressed genes compared to PB profiles. **d**. Volcano plots show robust DEGs between PB and bulk profiles, highlighting consistent transcriptomic differences. **e**. SOM portraits generated from the ovarian cancer dataset reveal distinct nonlinear expression topologies between PB and bulk, emphasizing modality-driven divergence. **f**. GO enrichment analysis shows PB-enriched DEGs are involved in conserved intracellular processes, while bulk-enriched DEGs reflect context-specific biological functions. SPCA – Sparse Principal Component Analysis; KDE – Kernel Density Estimate; DEG – Differentially Expressed Gene; GO – Gene Ontology; SOM – Self-Organizing Map.

Next, to quantify transcriptomic similarity, we computed Pearson correlations (r) across three comparison types: within bulk (bulk–bulk), within PB (PB–PB), and between bulk and PB (bulk– PB). In ovarian cancer, bulk–bulk correlations ranged from *r* = 0.89 to 0.96, and PB–PB correlations ranged from *r* = 0.83 to 0.98, reflecting strong within-modality consistency. In contrast, bulk–PB correlations were markedly lower, ranging from *r* = 0.76 to 0.89. Similar trends were observed in the breast cancer (**Figure 1b**) and iPSC (**Supplementary Figure 2b**) datasets, where bulk–PB correlations were consistently lower than within-modality comparisons. These findings reveal persistent divergence between PB and bulk RNA-seq profiles, suggesting that PB may not fully recapitulate true bulk expression landscapes. To further investigate this divergence at a distributional level, we compared the global transcriptomic distributions using kernel density estimates (KDE) of averaged normalized expression. Across all datasets, PB profiles showed a higher abundance of low-expression genes compared with bulk RNA-seq profiles (**Figure 1c; Supplementary Figure 2c**). This discrepancy likely reflects scRNA-seq-specific dropout effects, resulting in overrepresentation of low- abundance transcripts in PB profiles.

To further characterize fine-scale nonlinear transcriptomic differences, we applied Self-Organizing Map (SOM) analysis to cluster genes with similar expression dynamics, to observe and compare the global gene expression patterns between PB and bulk profiles. SOM analysis revealed two distinct patterns: (i) reciprocal expression clusters, where genes highly expressed in one modality were lowly expressed in the other, and (ii) non-overlapping modality-specific clusters unique to either PB or bulk profiles (**Supplementary Figure 3a**). In ovarian cancer, PB and bulk profiles clustered distinctly regardless of patient heterogeneity (**Figure 1e**). Similarly, in iPSC-derived cell lines, PB and bulk profiles differences persist despite genetic homogeneity (**Supplementary Figure 3b**); and in breast cancer, this separation was consistently observed across all clinical subtypes (**Supplementary Figure 3c**). These findings underscore that PB and bulk RNA-seq capture complementary rather than equivalent transcriptomic signals and reflect inherent biological properties of each profiling modality rather than mere technical artifacts.

To identify biological differences between PB and bulk RNA-seq, we employed an integrated approach combining classical differential expression (DE) analysis with multivariate feature selection. Using DESeq2 with standard thresholds (FDR < 0.05, |log2FC| > 1), we initially identified 7,998 differentially expressed genes (DEGs) in the ovarian cancer dataset. To further enhance the robustness of this candidate pool, we integrated SPCA results, noting that PC1 significantly discriminated samples by data modality (p < 0.001, t-test). By intersecting the DEGs with genes exhibiting non-zero loading in PC1, we obtained a refined set of 2,641 highly informative DEGs most strongly associated with differences between bulk and PB profiles (**Figure 1d**). Applying this refinement to the other datasets, we retained 3,808 of the 14,050 initial DEGs in breast cancer and 3,243 of the 6,467 DEGs in the iPSC-derived samples (**Supplementary Figure 2d**).

Subsequent Gene Ontology (GO) enrichment analysis of these refined DEGs revealed distinct functional signatures. PB-upregulated genes were consistently enriched for conserved intracellular processes like mitochondrial function and cytoplasmic translation, across all datasets (**Figure 1f**, right panel). Conversely, bulk-upregulated genes showed context-specific enrichments reflecting tissue-level biology. For example, in ovarian cancer, terms like epithelial sheet morphogenesis and suppressed macroautophagy suggested epithelial-mesenchymal transition (EMT) activation [11]. Similarly, breast cancer samples were associated with P-body assembly, regulation of small GTPase signaling, and chromatin-associated phosphorylation events, processes linked to breast cancer progression [12] (**Figure 1f**, left panel). In iPSC-derived samples, enriched terms included nucleolar chromatin organization, neural tube closure, brain development, and anoikis, which are processes associated with stem cell development [13, 14] (**Supplementary Figure 2e**). These findings highlight that while PB profiles capture core intracellular programs, bulk RNA-seq provides unique insights into emergent, tissue-level processes shaped by complex cellular interactions.

In conclusion, our findings raise important considerations regarding the assumption that PB can serve as a direct surrogate for bulk RNA-seq. While PB offers computational advantages, it consistently shows divergence from true bulk profiles across different case studies. These discrepancies can cause misleading interpretations in studies focused on biological processes such as cell–cell communication, extracellular matrix interactions, and other tissue-level dynamics attenuated in data derived from scRNA-seq. As digital twins and synthetic data generation using Generative AI are gaining momentum [15, 16], we caution against using PB and bulk RNA-seq interchangeably for benchmarking, model training and downstream analysis. To develop reliable biological models and robust translational insights, we strongly encourage future work to acknowledge and address these distinct characteristics of PB and bulk RNA-seq.

## Methods

### 1 Transcriptomics data overview and preprocessing

To systematically compare bulk RNA-seq and matched single-cell RNA-seq-derived pseudo-bulk (PB) profiles, we analyzed transcriptomic data from three independent studies, each containing paired bulk and single-cell RNA-seq datasets.

#### 1.1 Breast cancer with clinical subtypes (GSE176078)

The GSE176078 dataset includes matched bulk and scRNA-seq profiles from breast cancer patients, stratified into three clinical subtypes: HER2+ (n = 4), ER+ (n = 11), and triple-negative breast cancer (TNBC; n = 9). PB profiles were generated by aggregating raw scRNA-seq counts across all cells for each corresponding sample. Two samples lacking matched bulk RNA-seq (GSM5354532 and GSM5354533) were excluded. Genes expressed in fewer than 20% of both bulk and PB samples were filtered out, resulting in 23,191 genes for analysis. Count data were normalized using the trimmed mean of M-values (TMM) method implemented in the edgeR package [17]. Expression values were then converted to counts per million (CPM) via edgeR::cpm() and log-transformed using the natural logarithm with a pseudocount of 1 (log1p() in R [18]) to stabilize variance and adjust for library size differences.

### 1.2 High-grade serous ovarian cancer (GSE217517)

To validate trends observed in breast cancer, we incorporated the GSE217517 dataset, which includes matched bulk RNA-seq and scRNA-seq data from high-grade serous ovarian cancer patients. PB profiles were generated as described above. For patients with technical replicates in bulk RNA-seq, expression values were averaged due to high intra-replicate correlation (mean Pearson r = 0.95), yielding one representative bulk profile per patient. One sample (P8; GSM6720930 & GSM6720956) was excluded due to excessive missing data. Preprocessing steps followed the same pipeline as GSE176078, resulting in log-transformed CPM data for 20,112 genes.

#### 1.3 Induced pluripotent stem cell-derived lines (GSE226163)

The GSE226163 dataset includes biologically homogeneous samples consisting of endothelial and vascular smooth muscle cells differentiated from human induced pluripotent stem cells (iPSCs). Bulk (GSM7066034–GSM7066039) and scRNA-seq (GSM7066040–GSM7066045) data were available for each cell type, with three technical replicates per modality. PB profiles were constructed by aggregating gene expression across all scRNA-seq cells within each replicate, preserving one-to-one correspondence with bulk profiles, consistent with the structure of the other datasets. After applying the same filtering and normalization steps, 16,596 genes were retained for downstream analysis.

## 2 Comparison of bulk and PB RNA-seq data using data analytics and machine learning

### 2.1 Sparse principal component analysis (SPCA)

To assess global expression patterns of PB and bulk RNA-seq samples, we performed SPCA on the log-transformed CPM values using the *sparsepca*::spca() function [19] with centering and scaling enabled. Both *alpha* and *beta* parameters were set to 0.001, and the maximum iterations set to 500 for convergence. Ten principal components (PCs) were calculated, and we use the first two components, PC1 and PC2, which accounted for the largest proportion of variance, for visualizing sample distributions of PB and bulk profiles using the *ggplot2* [20] and *patchwork* [21] packages.

#### 2.2 Pearson Correlation Analysis

To quantify transcriptomic similarity, Pearson correlation coefficients were computed across three comparisons: within bulk (bulk-bulk), within PB (PB-PB), and between bulk and PB (bulk-PB) profiles using the cor() function in R. Results were visualized as boxplots using *ggplot2* and *patchwork* packages to quantify baseline differences across samples.

#### 2.3 Kernel Density Estimation (KDE)

To assess distributional shift, KDE was performed to the log-transformed CPM data using *ggplot2*::geom_density(). This visualization enabled qualitative assessment of modality-driven variation in gene expression distribution

#### 2.4 Self-Organising Maps (SOMs)

Nonlinear gene expression patterns were analyzed using the *oposSOM* R package [22]. Log-normalized expression profiles from all PB and bulk RNA-seq samples within each experiment were used to train SOMs with default parameters and a grid size of 40 × 40 nodes to achieve high-resolution clustering. Each SOM node represents a gene module comprising genes with coherent expression patterns across samples. Node-wise expression weights were visualized as heatmaps, providing biologically meaningful interpretation and cross-modality comparison of bulk and PB profiles.

#### 3 Biological signature divergence between PB and bulk RNA-seq

### 3.1 Differential gene expression (DGE) analysis

Differential gene expression analysis was performed using DESeq2 [23] with raw gene counts as input to identify significantly differentially expressed genes (DEGs) between PB and bulk RNA-seq profiles. Genes with an absolute log_2_ fold change > 1 and a false discovery rate (FDR) < 0.05 were considered significant.

To enhance the robustness of DEG selection, we integrated results from Sparse Principal Component Analysis (SPCA). Among the top principal components (PCs), only PC1 consistently showed significant separation between PB and bulk profiles (p < 0.0001) and was therefore used for downstream filtering. Genes with non-zero loadings in PC1 were retained for further analysis.

To infer modality-specific associations, we compared the average PC1 scores between PB and bulk RNA-seq samples. If bulk samples exhibited lower or negative average PC1 scores, genes with negative loadings were considered to contribute predominantly to the bulk RNA-seq signature, while genes with positive loadings were associated with PB profiles. This interpretation was reversed if PB samples showed lower average PC1 scores. The final set of informative DEGs was defined as the intersection of SPCA-selected genes (from PC1) and DESeq2-derived DEGs, representing modality-specific transcriptional signatures. Volcano plots were generated to visualize log_2_ fold changes and FDR values, highlighting these informative DEGs.

### 3.2 Gene Ontology (GO) analysis

GO enrichment analysis was conducted using the topGO package [24] with informative DEGs as input and all expressed genes per sample as the background gene set. The weight01 algorithm was used to account for GO graph topology, and significance was assessed using Fisher’s exact test. GO terms were ranked by p-value within each sample, and the top 5 most significantly enriched terms of each sample were selected and visualized by gene ratio and significance for each modality.

## Data availability

All data used in our study can be found on Gene Expression Omnibus (GEO). Ovarian cancer dataset is found under accession GSE217517, breast cancer dataset with accession GSE176078 and iPSC dataset with accession GSE226163.

## Code availability

Source code for all analysis scripts and pipelines is available at GitHub (https://github.com/boonyhow/BulkPB-analysis)

## Online content

Supplementary figures are available at Brief Communication online.

## Author contributions

B.H.L. developed the computational pipelines for transcriptomic analyses. B.H.L. and M.M.R. jointly generated all visualizations and co-wrote the manuscript. K.S. conceptualized the study, supervised and edited the manuscript.

