## Supplementary Figure 1 for "Machine learning differentiates between bulk and pseudo-bulk RNA-seq datasets"

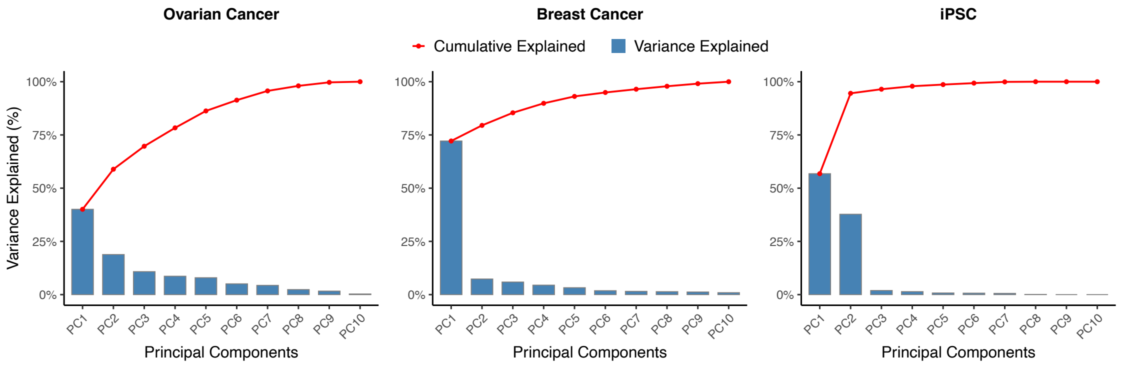


**Supplementary Figure 1**. The proportion of variance explained by Sparse Principal Component Analysis (SPCA) components indicates that PC1 accounts for the largest variance across all datasets, including ovarian cancer, breast cancer, and induced pluripotent stem cell (iPSC) lines. This suggests that modality-driven differences between bulk and pseudo-bulk RNA-seq profiles dominate over other sources of variation
