## Supplementary Figure 2 for "Machine learning differentiates between bulk and pseudo-bulk RNA-seq datasets"

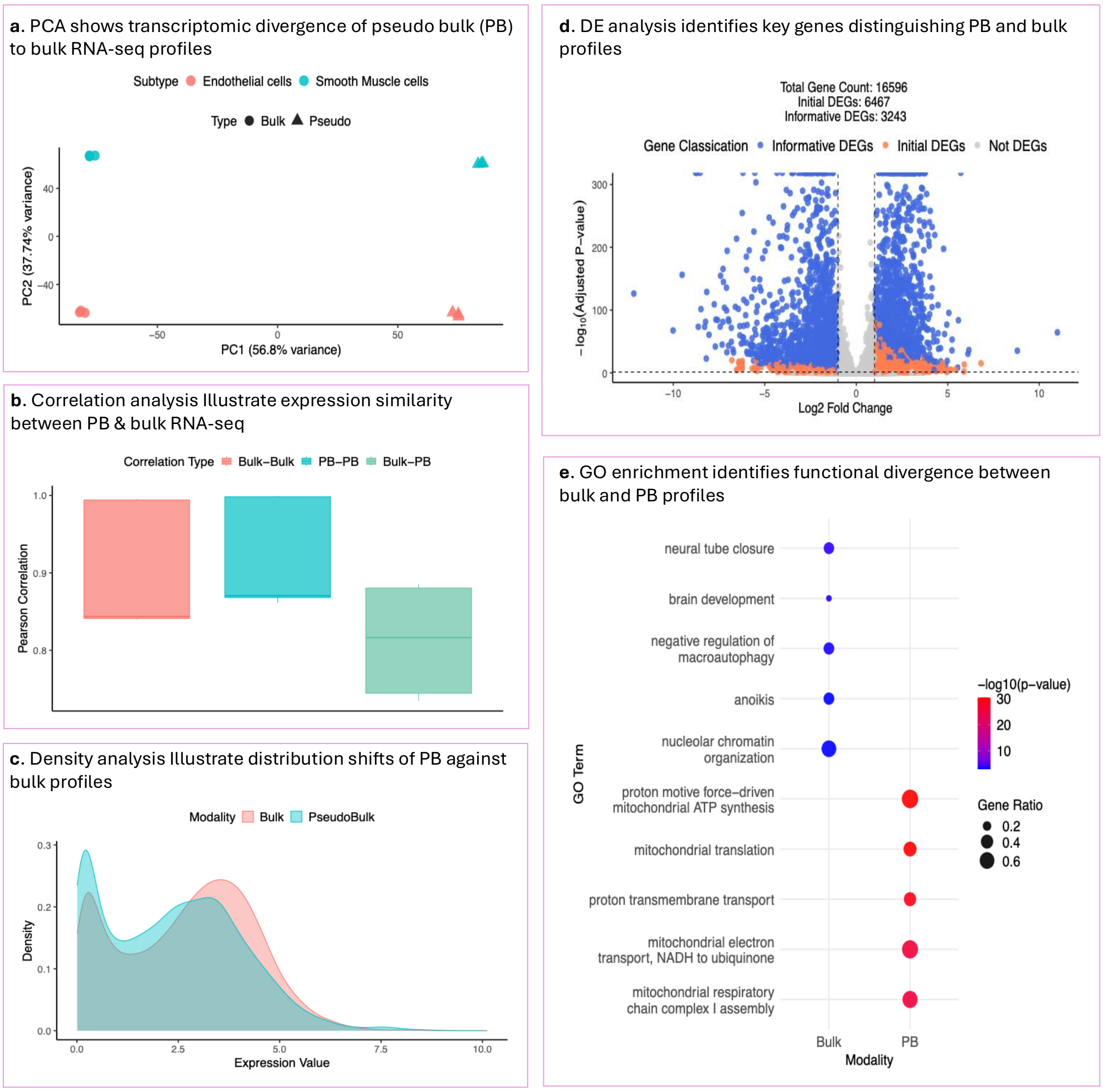


**Supplementary Figure 2. Transcriptomics divergence between PB and bulk RNA-seq profiles observed in homogeneous iPSC-differentiated cell lines.**

**a.** SPCA shows that PC1 distinctly separates samples by data modality, whereas PC2 captures variation across cell line identities. **b.** Pearson correlation analysis shows consistently high within-modality similarity, whereas similarity between PB and bulk samples are markedly lower, consistent with trends observed in breast cancer and ovarian cancer datasets. **c.** Density analysis reveals right-shifted expression distribution in bulk samples, illustrating a greater proportion of highly expressed genes relative to PB profiles **d.** Volcano plots show robust DEGs between PB and bulk profiles. **e.** GO enrichment analysis shows PB-enriched DEGs are involved in intracellular processes, aligning with patterns observed in breast cancer and ovarian cancer, while bulk-enriched DEGs reflect biological functions associated with stem cell differentiation. SPCA: Sparse Principal Component Analysis; KDE: Kernel Density Estimate; DEG: Differentially Expressed Gene; GO: Gene Ontology; SOM: Self-Organizing Map.
