## Supplementary Figure 3 for "Machine learning differentiates between bulk and pseudo-bulk RNA-seq datasets"

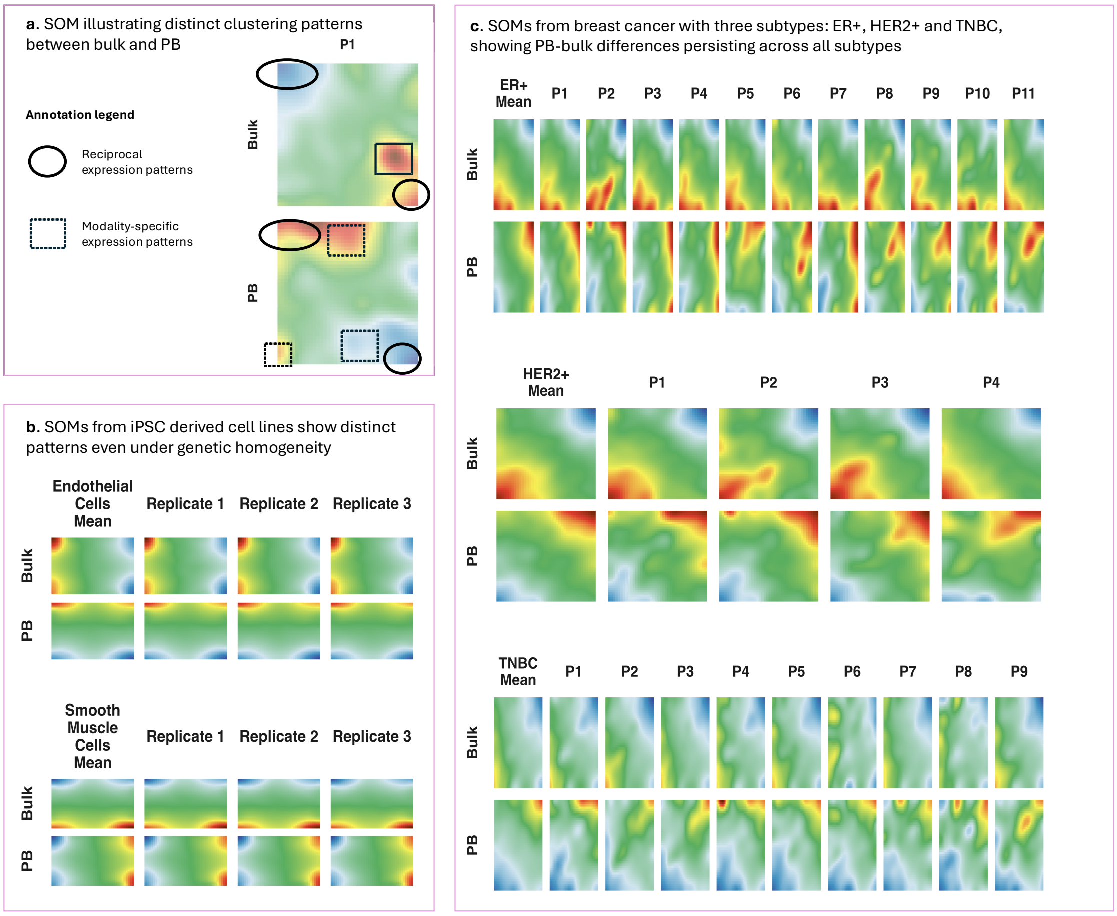


**Supplementary Figure 3. SOMs consistently highlight nonlinear transcriptomic differences between PB and bulk profiles across datasets.**

**a.** SOMs visualization of ovarian cancer sample P1 (from main text **Figure 1e**) shows annotated expression regions. Circles highlight reciprocal modality-divergent regions, where high expression in one modality corresponds to low expression in the other. Squares indicate modality-specific high-expression clusters, reflecting distinct transcriptomic contributions unique to PB or bulk RNA-seq. **b.** SOMs of iPSC-derived cell types (Endothelial and Smooth Muscle Cells) show consistent clustering across biological replicates, reflecting high genetic homogeneity. Nevertheless, distinct expression patterns are observed between PB and bulk profiles from the same replicate, suggesting that modality-driven differences persist even in homogeneous cellular contexts. **c.** SOMs of breast cancer patients stratified by clinical subtype (ER+, HER2+, TNBC) show that PB–bulk differences are consistently preserved across all subtypes. ER+ and HER2+ patients display similar expression patterns across individuals, but TNBC shows slightly different clustering patterns, suggesting greater heterogeneity. Nonetheless, PB and bulk profiles remain clearly distinguishable within each patient, emphasizing the modality-dependent biological differences captured by the two RNA-seq approaches
